# Design of a Water-Soluble CD20 Antigen with Computational Epitope Scaffolding

**DOI:** 10.1101/2024.12.05.627087

**Authors:** Zhiyuan Yao, Brian Kuhlman

## Abstract

The poor solubility of integral membrane proteins in water frequently hinders studies with these proteins, presenting challenges for structure determination and binding screens. For instance, the transmembrane protein CD20, which is an important target for treating B-cell malignancies, is not soluble in water and cannot be easily screened against potential protein binders with techniques like phage display or yeast display. Here, we use *de novo* protein design to create a water-soluble mimic of the CD20 dimer (“soluble CD20”). Soluble CD20 replaces the central transmembrane helix of CD20 with a water-soluble helix that dimerizes to form a coiled coil that structurally matches the dimer interface of native CD20 and presents the central extracellular loop of CD20 in a binding competent conformation. Unlike peptides derived from CD20, soluble CD20 binds tightly to monoclonal antibodies that recognize quaternary epitopes on the extracellular face of CD20. We demonstrate that soluble CD20 is easy to produce, remains folded above 60°C, and is compatible with binder screening via yeast display. Our results highlight the ability of computational protein design to scaffold conformational epitopes from membrane proteins for use in binding and protein engineering studies.

**Importance:** This study demonstrates that computational protein design can be used to create water soluble variants of transmembrane proteins. The water-soluble CD20 mimic created here is compatible with high throughput binding screens and should be useful for developing next generation anti-CD20 therapeutics.

## Introduction

Therapeutic antibodies targeting multi-spanning membrane proteins (or complex membrane proteins, CMPs) remain underdeveloped despite their importance in human disease. While antibody therapeutics provide exceptional specificity and affinity, making them a transformative class of therapeutics, their development has largely focused on soluble ligands or transmembrane receptors with large extracellular domains (ECDs) (Stephens and Wilkinson 2024; Crescioli et al. 2024). Despite CMPs comprising approximately 60% of small-molecule drug targets, only 20 among 221 approved therapeutic antibody therapeutics are targeting CMPs, including 13 targeting CD20 (Lim et al. 2010; Casan et al. 2018), 4 targeting CGRP (Aditya and Rattan 2023; Cohen et al. 2022), 1 targeting GPCR5d (Rodriguez-Otero et al. 2024), 1 targeting CCR4 (Zengarini et al. 2024), and 1 targeting Claudin 18.2 (M. A. Shah et al. 2023).

Developing antibodies or other protein binders against CMPs is hindered by challenges associated with producing well-folded CMPs (To’a Salazar et al. 2021; R. Dodd et al. 2020; Hashimoto et al. 2018). CMPs are membrane proteins that span cell membranes with multiple hydrophobic transmembrane domains (Qing et al. 2022). These proteins are critical drug targets due to their role in cell signaling, immune responses, and various human diseases (To’a Salazar et al. 2021; Stephens and Wilkinson 2024; R. B. Dodd, Wilkinson, and Schofield 2018). Access to soluble, well-behaved, conformation correct antigens is important for efficient binder development by animal immunization or high-throughput screening technologies like phage or yeast display. Due to their hydrophobic surfaces, CMPs have low solubility and are often unstable when isolated from the lipid bilayer environment (Qing et al. 2022). Several approaches have been developed to solubilize CMPs (Krishnarjuna and Ramamoorthy 2022), including the use of detergent micelles (Cecchetti et al. 2021; Kotov et al. 2019), proteoliposomes (Yao, Fan, and Yan 2020; Takeda et al. 2015), nanodiscs (Frauenfeld et al. 2016; Ma et al. 2020), or virus-like particles (Tucker et al. 2018). These approaches typically require extensive optimization to balance solubility and protein stability, making it both time-consuming and expensive. An emerging alternative is the creation of water-soluble analogs of MPs. By redesigning hydrophobic transmembrane domains to increase hydrophilicity while preserving selected epitopes from the MP, these analogs enable binder development with high-throughput screening platforms (Qing et al. 2022). Several previous studies systematically replaced hydrophobic amino acids with hydrophilic residues of similar size and shape (Zhang et al. 2018; Smorodina et al. 2022), and successfully generated solubilized GPCRs (Skuhersky et al. 2021), ion channels (Smorodina et al. 2024), and chemokine receptors (Qing et al. 2023). Recently, AI-based methods for protein design have been used to generate soluble proteins that present selected structural motifs from CMPs (Goverde et al. 2024).

Here, we use computational protein design to create a soluble version of the multi-spanning membrane protein CD20 for use as “bait” in binder discovery campaigns. CD20 is a CMP expressed on B cells and is an effective therapeutic target in the treatment of B-cell malignancies (McLaughlin et al. 1998; Luo et al. 2021). Over the last 20 years, 13 different anti-CD20 antibodies have been developed and approved (Cragg and Glennie 2004; Alduaij and Illidge 2011; Klein et al. 2013). Because native CD20 is insoluble in water, these antibodies were generated through animal immunization with CD20-overexpressing cells as opposed to immunization with soluble recombinant protein. This approach has worked well for generating antibody binders, but it is incompatible with high-throughput technologies such as yeast (Cherf and Cochran 2015) and phage display (Jaroszewicz et al. 2022). These advanced methods not only facilitate the optimization of antibodies derived from animal immunization, but also enable the discovery of non-antibody binders, such as miniproteins (Cao et al. 2022) and DARPins (Plückthun 2015) that can be exceptionally stable (Kortemme 2024) and can be linked to create multi-specific fusion proteins (Huang et al. 2024). The enhanced ability to discover novel CD20 binders would meet the emerging demand for developing next-generation CD20 therapeutics (Dabkowska, Domka, and Firczuk 2024), which focus on multispecific targeting to combat resistance (Beers et al. 2010; Shah et al. 2020; Fousek et al. 2021) and refining well-folded antigen recognition domains essential for chimeric antigen receptor (CAR) therapies (Chen et al. 2023; Zhen et al. 2024).

To engineer a water-soluble version of CD20, we used computational de novo protein design to create a protein scaffold that presents the primary extracellular loop of CD20 in a native-like conformation. Unlike the solubilized native CD20 in detergent micelles, our engineered soluble CD20-mimic is compatible with screening via yeast display. We demonstrate that the CD20 mimic expresses well, is thermostable, and can bind both type I and type II anti-CD20 antibodies with native like affinities and stoichiometries. Our results demonstrate how computational protein design can be used to solubilize transmembrane proteins and the designed CD20 mimic should be a powerful reagent for discovering next-generation anti-CD20 therapeutics.

## Results

### Design of a soluble CD20 mimic

The CD20 monomer consists of four transmembrane α-helices (TM). The N- and C-termini are located on the intracellular side of the membrane and extracellular loops connect TM1 with TM2 and TM3 with TM4 (Figure 1)(Kumar et al. 2020; Rougé et al. 2020). The first extracellular loop (ECL1) is short and largely buried by the second extracellular loop (ECL2). In the membrane, CD20 forms a homodimer with the majority of the interchain contacts coming from ECL2, TM4 and TM1. On the extracellular face of the homodimer, a cleft is formed between the two copies of ECL2. This cleft and the surrounding residues from ECL2 is a binding site for some anti-CD20 antibodies(Kumar et al. 2020; Rougé et al. 2020). We hypothesized that a de novo designed homodimer that presents residues from ECL2 in the correct orientation relative to each other and forms a native-like cleft would be an effective mimic of the extracellular binding surface of CD20.

**Figure 1.**
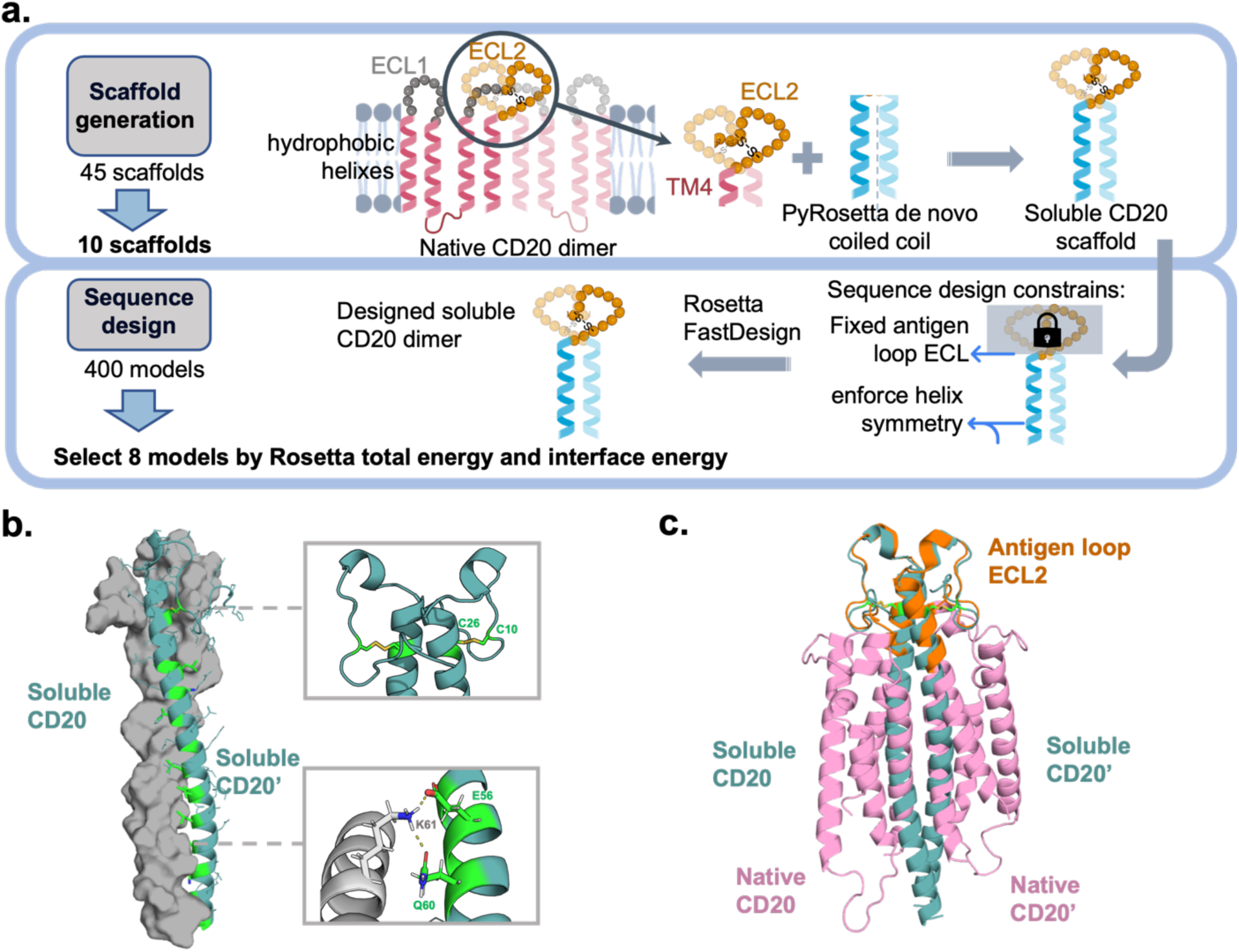
Design of a water soluble CD20 dimer mimic using Rosetta Design. a) The ECL2 loop of CD20 was grafted onto dimeric coiled coils and regions outside the antigen loop were sequence optimized with the Rosetta FastDesign protocol (Maguire et al. 2021). b) Final design model of soluble CD20 with one chain shown as a gray surface and one chain represented in ribbon mode. The native disulfide from the ECL2 loop is conserved in the design (top right inset) and the coiled coil dimer is stabilized by hydrophobic interactions and hydrogen bonding (lower right inset). c) A superposition of soluble CD20 (blue) and native CD20 (pink).

The first several residues of TM4 sit above the membrane, serve as an anchor for the C-terminal portion ECL2, and make contacts with TM4 from the partner chain. To stabilize the binding cleft formed by the ECL2/ECL2 interface in a soluble protein, we sought to scaffold ECL2 and the N-terminal portion of TM4 with a de novo designed coiled coil. First, we generated a large family of idealized dimeric coiled-coils by varying the standard Crick parameters for a coiled-coil, which describe the superhelical twist and radius of a coiled-coil (Grigoryan and DeGrado 2011a). Ten coiled coils were selected, and ECL2 of CD20 cryo-EM structure were grafted onto the N-terminus of the matching coiled coils to construct full-length models of the designed proteins, each containing sixty-eight residues.

Sequences were designed for the CD20/coiled coil chimeras using the FastDesign protocol in the molecular modeling program Rosetta. FastDesign iterates between rotamer-based sequence optimization and backbone refinement to find low energy sequence/structure pairs (Maguire et al. 2021). The CD20/coiled coil chimeras were designed to form a homodimeric structure, with the antigen epitope loop (ECL2) preserved in its native sequence and conformation, including conservation of the native disulfide between residues 167 and 183 (Figure 1, b). The dimer interface of the designed coiled coil is predominantly stabilized by hydrophobic interactions, forming a tightly packed core that ensures structural stability. In addition to these hydrophobic contacts, a few key hydrogen bonds are observed, contributing to the specificity and further reinforcing the dimerization. When aligned with the native CD20 dimer cryo-EM structure, the designed chimera preserves the ECL2 antigen loop in its native conformation, although the coiled coil regions differ significantly from the native transmembrane helices TM1 and TM4 (Figure 1, c). Eight out of 400 models were selected for experimental validation.

### Soluble CD20 forms a stable dimer

The 8 selected design sequences were fused to a N-terminal signal sequence and a C-terminal region that included a FLAG tag, His-tag, and Avitag and were expressed as secreted proteins in Expi293 cells. Following purification with metal affinity chromatography, two of the design sequences produced detectable amounts of protein (Figure S1), with the highest yields observed for design F2 (33.1 mg per liter of cell culture). Design F2 was renamed “soluble CD20”. In the following experiments we make use of three different versions of the soluble CD20 protein. Soluble CD20^Avi^ contains a C-terminal FLAG-tag, His-tag and Avi-tag and was the construct used in the initial screening.

Soluble CD20^His^ includes a N-terminal His-tag followed by a TEV protease site, and soluble CD20^bacteria^ was constructed in a bacterial expression vector (full sequences shown in supplementary table 2). We tested the expression of soluble CD20^bacteria^ in E. coli. using Shuffle T7 cells. Final yields after purification were 4 mg per liter of cell culture. In contrast to soluble CD20, native CD20 cannot be secreted as a soluble protein from Expi293 cells and yields were low (0.44 mg/L) when expressed in SF9 cells and purified by membrane solubilization (Rougé et al. 2020) (Figure S1,S2).

On a reducing SDS-page gel, soluble CD20^Avi^ runs with an apparent molecular weight of 15 kDa, close to the expected molecular weight (14.9 kDa) of dissociated monomer (Figure 2a). To determine the oligomerization state of soluble CD20 in solution, size-exclusion chromatography with multi-angle light scattering (SEC-MALS) was performed. Soluble CD20^Avi^ eluted as a single peak with a molecular weight of 29.0 ± 0.9 kDa (Figure 2b), consistent with the expected molecular weight for a homodimer 29.8 kDa. Soluble CD20 has a circular dichroism (CD) spectrum consistent with a helical protein and unfolds cooperatively with temperature (Figure 2c). As expected for a homodimer, the thermal stability of soluble CD20 is concentration dependent with a thermal unfolding midpoint of 68.6°C at a protein concentration of 2 μM (Figure 2d). Overall, biochemical characterization confirmed that the soluble CD20 design was expressed at the expected molecular weight and assembled into a stable dimer.

**Figure 2.**
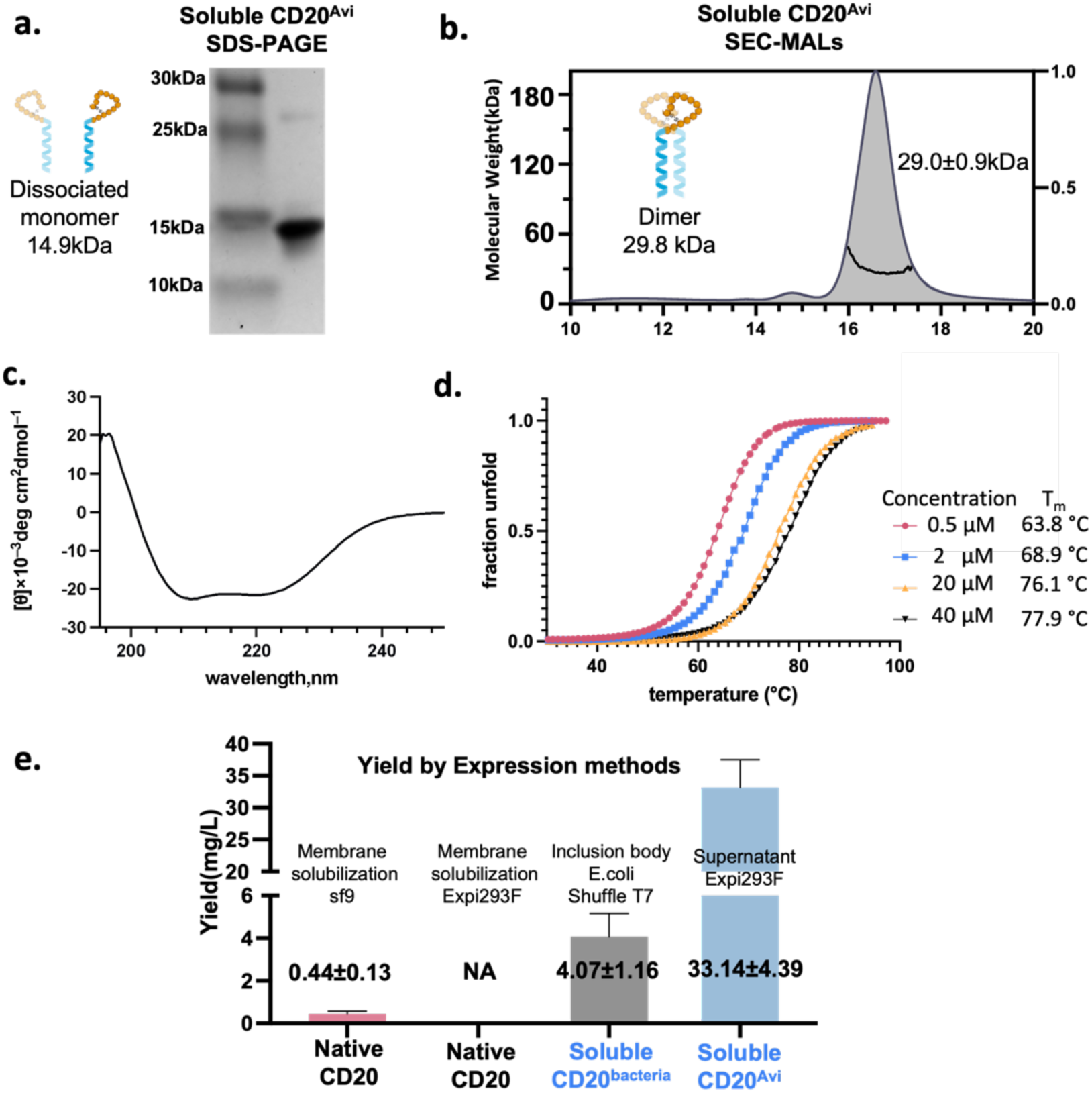
Expression and biophysical characterization of soluble CD20 as a stable dimer. a) Purified CD20^Avi^ visualized with SDS-PAGE. b) SEC-MALS profile of soluble CD20 shows a single peak around 29 kDa, which corresponds to its expected molecular weight of 29.8 kDa as a dimer. c) Circular dichroism spectra and d) thermal melts™ of CD20^Avi^ at the indicated protein concentrations. CD data suggest that De novo CD20 adopts an α-helical secondary structure Tm increases in higher concentration suggests De novo CD20 assembles as oligomer and dissociates when temperatures increase. e) Expression yields of selected native and soluble CD20 constructs from sf9, Expi293F, or E. coli cells. Native CD20 is unattainable from Expi293 supernatant. While de novo CD20 achieves expression yield at 33mg/L.

### Soluble CD20 design binds to both type I and type II antibodies with native-like stoichiometry

Previously discovered anti-CD20 mAbs have been categorized into two types: type I, ‘rituximab(RTX)-like’ mAbs, which induce complement-dependent cytoxicity (CDC) upon binding by crosslinking CD20 receptor, and type II anti-CD20 mAbs, such as obinutuzumab(OBZ), which do not induce accumulation of CD20 and show relatively little CDC activity (Klein et al. 2013). Cryo-EM structures of the fragment antigen-binding region (Fab) of type I and II mAbs bound to the native CD20 dimer show different stoichiometries and binding geometries (Figure 3a)(Kumar et al. 2020; Rougé et al. 2020). The type I antibody rituximab binds CD20 with two Fab molecules per CD20 dimer. The complex is stabilized by homotypic Fab-Fab interactions which contribute to a slow off rate of rituximab from the CD20 dimer. In contrast, the type II mAb OBZ binds CD20 with one Fab molecule per CD20 dimer. These varying stoichiometries have been linked to the different functional properties of type I and II anti-CD20 mAbs.

**Figure 3.**
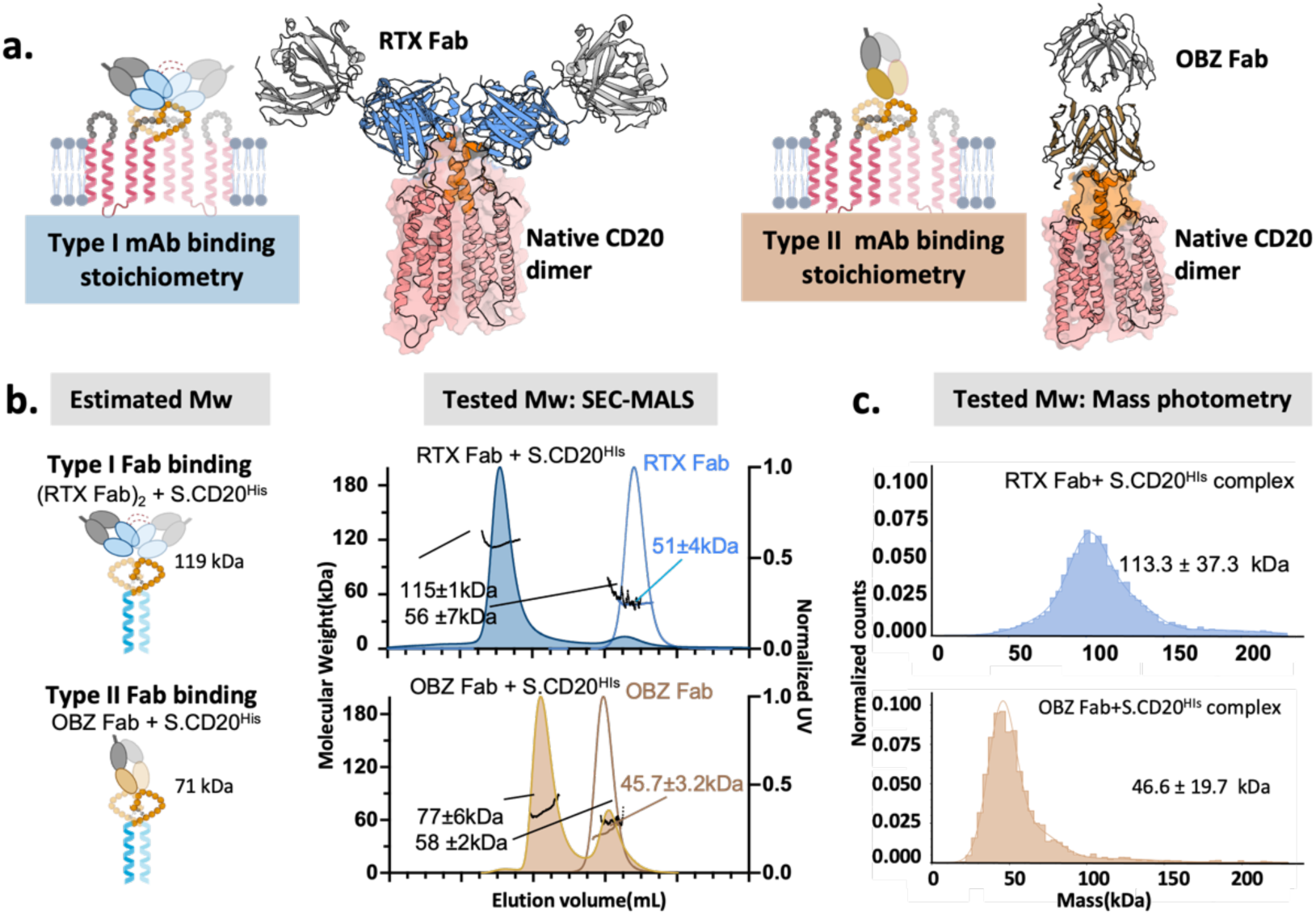
Soluble CD20 mimicked the binding stoichiometry of type I and type II antibodies a) Rituximab (RTX) is a type I anti-CD20 antibody that binds native CD20 with 2:2 stoichiometry. Obinutuzumab (OBZ) is a type II anti-CD20 antibody that binds native CD20 with 1:2 (Fab:CD20) stoichiometry. b) RTX and OBZ form complexes with soluble CD20^His^ and the measured molecular weights via SEC-MALS match the expected binding stoichiometries. c) Mass photometry experiments with soluble CD20^His^ mixed with RTX or OBZ. Protein concentrations were 50 nM.

To test if soluble CD20 closely mimics the extracellular epitopes of native CD20, we determined the binding stoichiometries of RTX and OBZ to soluble CD20. Fabs were mixed with soluble CD20 in a molar ratio of 2:2.5 and analyzed with SEC-MALS. With the RTX Fab, a large peak was observed with a measured molecular weight of 115 kDa which is close to the expected molecular weight (119 kDa) for a complex of two Fabs per soluble CD20^His^ dimer (2:2 stoichiometry). With the OBZ Fab, a peak was observed with a measured molecular weight of 77 kDa which is close to the expected molecular weight (71 kDa) for complex of 1 Fab per soluble CD20^His^ dimer, consistent with the 1:2 stoichiometry (Figure 3b). In the RTX experiment, a separate peak was observed for excess amount of soluble CD20 (Figure S3). In the OBZ Fab experiments, a secondary peak was observed for excess Fab. These results indicate that rituximab and OBZ bind soluble CD20 in a manner that closely mimics binding to native CD20.

To further interrogate antibody binding to soluble CD20, gel filtration was used to purify the soluble CD20/Fab complexes, and the samples were examined with mass photometry. The RTX sample displayed a single species in solution with a molecular weight of 113 kDa, consistent with a complex of two Fabs per molecule of soluble CD20^His^ dimer. In contrast, the OBZ sample only displayed a peak at 47 kDa, which aligned with the expected mass of unbound Fab, suggesting that OBZ could be dissociated from soluble CD20^His^ at the experimental conditions used to perform mass photometry. (Fig 3c) Mass photometry requires low protein concentrations for detection and cannot detect species below 30-40 kDa. These experiments were performed at a complex concentration of approximately 50 nM, which is lower than the dissociation constant of OBZ for soluble CD20. Under these conditions, partial dissociation of the soluble CD20/OBZ Fab complex may have occurred, resulting in the observation of a lower molecular weight Fab peak and unbound soluble CD20 unobserved. The mass photometry results demonstrate that soluble CD20 can differentiate between the binding stoichiometries of type I and type II anti-CD20 antibodies, which consistent with finding by SEC-MALS analysis. This suggests that soluble CD20 accurately replicates the stoichiometric behavior of native CD20 with these two different types of antibodies.

### Soluble CD20 binds with native-like affinities to type I and II antibodies

The equilibrium dissociation constants (K_D_) between soluble CD20^Avi^ and anti-CD20 mAbs were measured with bio-layer interferometry (BLI) (Figure 4). In these experiments, native CD20, CD20 antigen peptide, or soluble CD20^Avi^ were immobilized on the sensor tips and dipped into solution with varying concentrations of mAbs in an IgG or Fab format. Soluble CD20^Avi^ bound to RTX IgG with an affinity (1.6 nM) that is close to our measured affinity between native CD20 and RTX IgG (1.8 nM) as well as the reported affinity for this interaction (1.7 nM)(Rougé et al. 2020). In contrast, a commonly used antigen peptide derived from the ECL2 loop of CD20 only binds to RTX with an affinity of 2.2 μM. Soluble CD20^Avi^ displayed affinities for RTX Fab (K_D_ = 20 nM) and OBZ Fab (68 nM) that are very similar to previously reported K_D_s between native CD20 and RTX Fab (21 nM) and OBZ Fab (59 nM). We also observed that RTX Fab dissociates more slowly from soluble CD20^Avi^ than OBZ, consistent with the homotypic interactions formed between RTX Fabs when two Fab molecules bind to the CD20 dimer. Overall, these results indicate that soluble CD20 presents the epitopes for type I and type II antibodies in a manner that closely resembles native CD20.

**Fig. 4.**
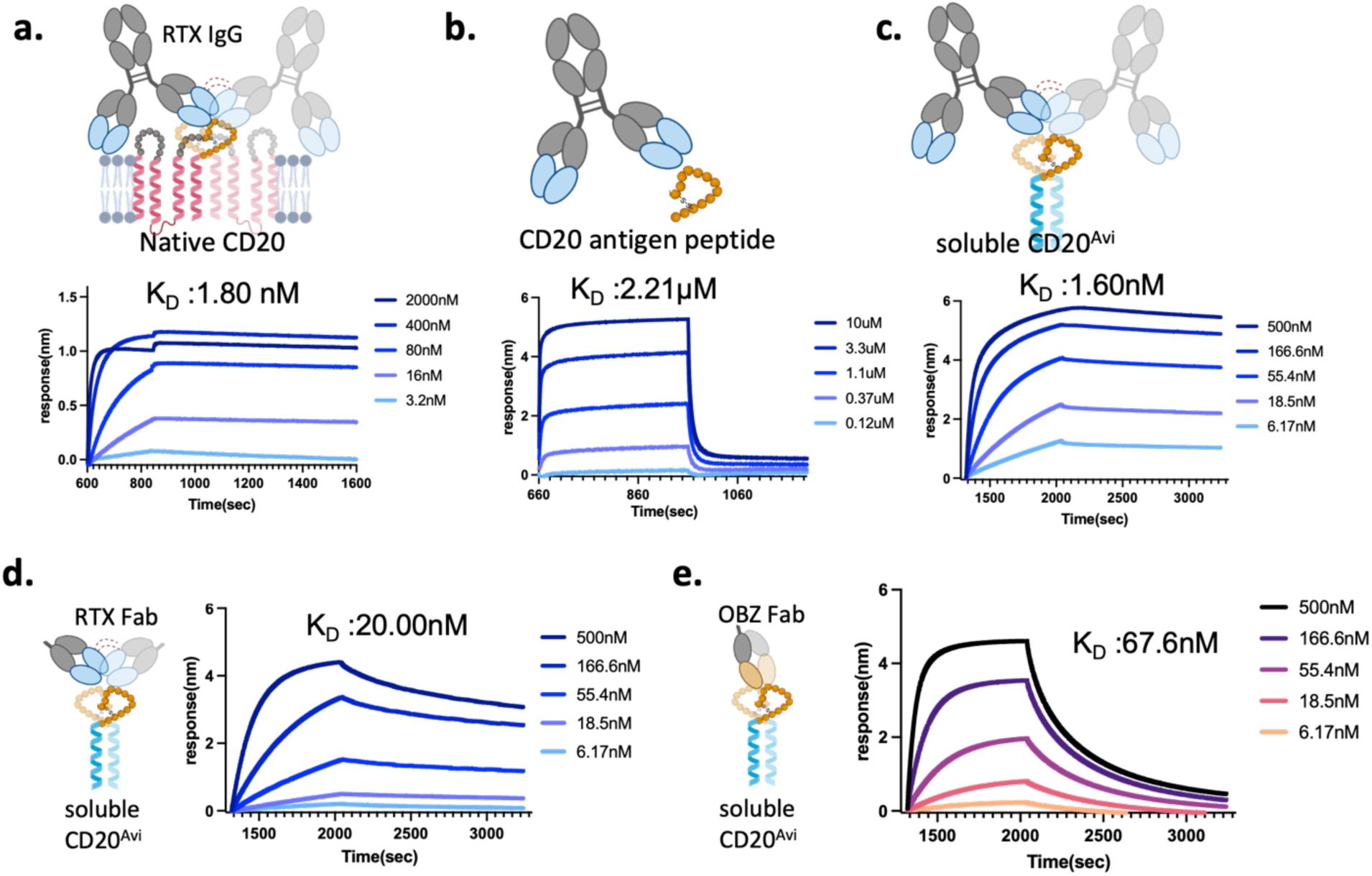
Soluble CD20 mimics the binding affinity of types I and II antibodies. BLI experiments with a) native CD20, b) CD20 peptide and c) soluble CD20^Avi^ immobilized on the BLI sensor tips and incubated with the indicated concentrations of RTX IgG. BLI experiments with soluble CD20^Avi^ binding to Fabs from d) RTX or e) OBZ. The measured binding affinity of soluble CD20 to RTX and OBZ is 20.00nM and 67.6, which is in proximity to native CD20 binding affinity 21.4nM and 58.8nM reported in literature.(Rougé et al. 2020; Kumar et al. 2020)

**Figure 5.**
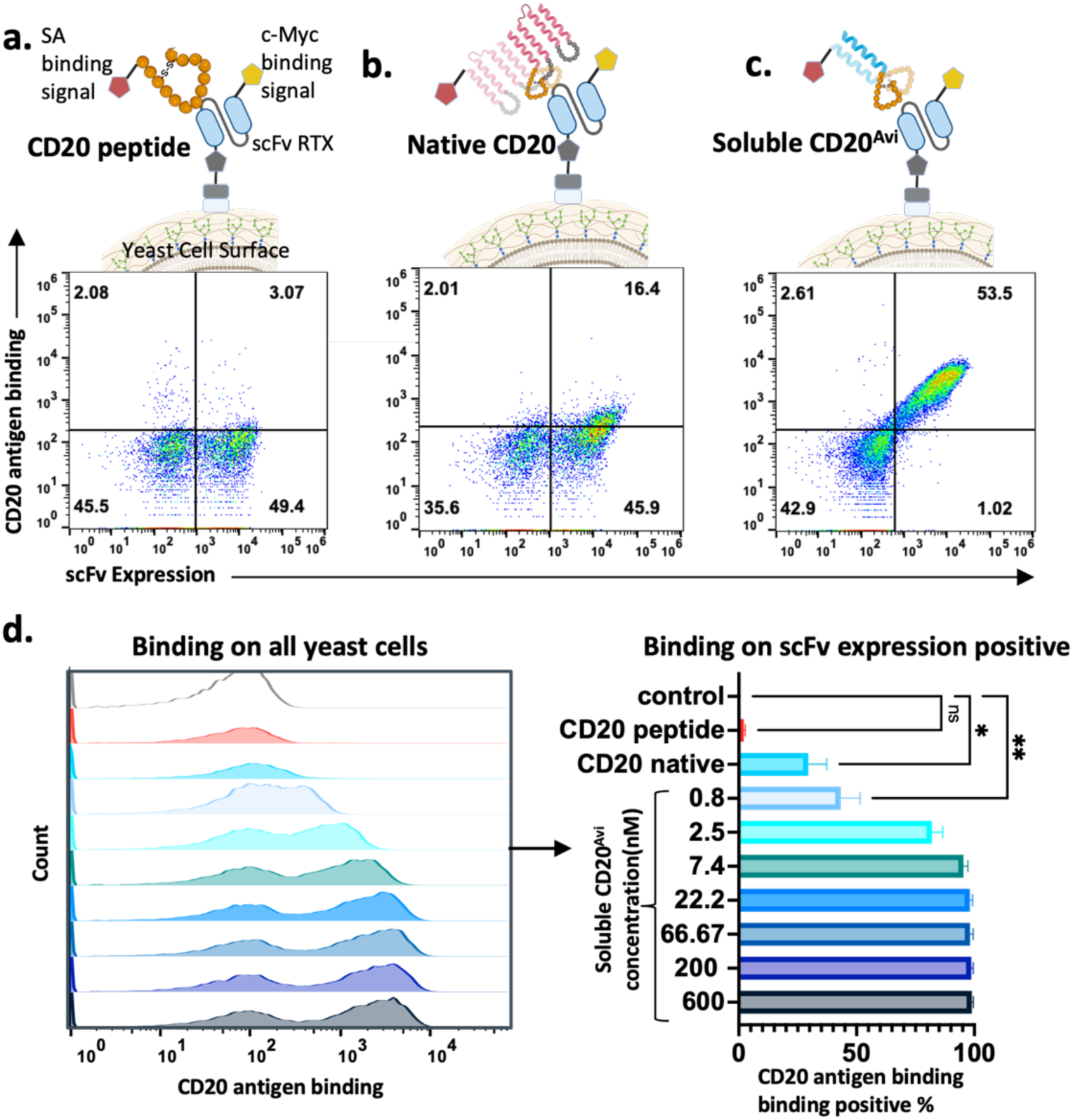
Soluble CD20 as a high-affinity antigen in yeast surface display. a,b,c) Yeast cells expressing a single-chain variable fragment (scFv) antibody derived from rituximab were tagged with a myc-tag to indicate expression levels. CD20 antigen forms — biotinylated CD20 peptide, native CD20, and soluble CD20 — were loaded onto yeast at concentrations of 10 µM, 0.6 µM, and 22.2nM, respectively. Binding was detected using fluorescently tagged streptavidin (SA). d) Concentration-dependent binding of soluble CD20 on yeast surface display. Binding-positive ratios represent the fraction of yeast cells with a positive SA binding signal among those positive for scFv expression. Control yeast cells, incubated with fluorescent SA alone, served to establish background fluorescence levels.

### Soluble CD20 can be used as a high-affinity antigen for yeast surface display

High-throughput display technologies such as phage, yeast, and mammalian display are efficient approaches for screening therapeutic binder libraries. These technologies typically require that the target antigen be stable, pure, and soluble to perform selection, presenting a significant challenge when dealing with membrane proteins. To test if engineered soluble CD20 is an effective target for yeast display experiments, we evaluated native CD20, CD20 primary epitope peptides, and our designed soluble CD20^Avi^ as target antigens in yeast surface display. We quantified the binding efficiency of these target proteins using yeast cells engineered to display a single-chain variable fragment (scFv) derived from rituximab. The expression level of the scFv on the yeast surface was measured using a C-myc tag at the scFv’s C-terminus, and the binding interactions between the displayed scFv and the target proteins were assessed via a fluorescent-labeled streptavidin signal, which binds to the biotinylated target protein(Chao et al. 2006).

The results indicate that a CD20 primary epitope peptide derived from ECL2 of CD20 is a poor target as low binding was detected to the rituximab scFv even at a peptide concentration of 10 µM. Native CD20 also exhibited low binding efficiency as a target antigen in the yeast display setup. When used as a soluble target protein, native CD20 required solubilization in detergent micelles, which may mask extracellular loops and non-specifically interfere with antibody binding. Despite high binding affinities characterized in biolayer interferometry (BLI) and surface plasmon resonance (SPR), native CD20 solubilized in surfactant micelles was inefficient as an antigen protein in yeast display at a concentration of 0.6 µM, with only 29.5 ± 7.8% of yeast cells displaying a positive binding signal. This aligns with the known challenges of using membrane proteins solubilized by detergent micelles in display technologies.

In contrast, the designed soluble CD20, which retained most antigenic features without the need for surfactant micelles, exhibited high binding efficiency as a target protein in yeast display. Among yeast cells expressing positive scFv, 98.1 ± 1.1% showed a positive binding signal at a soluble CD20 concentration of 22 nM. Further experiments were conducted with soluble CD20 concentrations ranging from 600 nM to 0.8 nM in 1:3 titrations to assess the binding efficiency on rituximab scFv-displayed yeast cells. Remarkably, 43.1 ± 8.2% of cells demonstrated binding at the lowest tested concentration of 0.8 nM. These findings underscore the efficacy of soluble CD20 as a high-performance target protein in yeast display settings.

## Discussion

There is long standing interest in methods that can be used to redesign complex membrane proteins (CMPs) to be folded and soluble in water (Rawlings 2016). One approach is to redesign hydrophobic amino acids on the surface of the transmembrane segments of the protein to be polar amino acids (Roosild and Choe 2005; Smorodina et al. 2024). Recent AI-based methods for sequence design have made this strategy more robust and have been used to solubilize GPCRs, the claudin-like fold, and a 6 transmembrane protein with the rhomboid protease fold (Goverde et al. 2024). One attractive aspect of this approach is that the protein remains full-length and if the design process is successful the overall fold of the protein should match its structure when embedded in the membrane. However, there are cases where it may only be important to maintain key structural features from the extracellular (or intracellular) components of the protein. In our study, our aim was to present the primary extracellular loop of CD20 as a homodimer with interchain positioning that mimics the wild-type protein. We were able to achieve this goal by scaffolding the loop with a *de novo* designed coiled-coil. Useful aspects of this approach include the high stability and solubility of de novo designed proteins and the ability to cut away regions of the membrane protein what may be problematic for downstream studies (Goverde et al. 2024). Also, this is a design approach that will benefit significantly from recent advances in motif-scaffolding with generative AI methods such as RFdiffusion (Watson et al. 2023).

Since full length CD20 is challenging to produce and is not easily adopted to screening or immunization studies, it has been common to use peptides and peptide fusion derived from the extracellular loops of CD20 as surrogates for the full-length protein. It has been hypothesized that peptides of this type could be used to elicit an anti-CD20 antibody response and provide active immunotherapy against autoimmune diseases (Perosa et al. 2005; Li et al. 2006; Liu et al. 2016). Our results highlight that these peptides are unlikely to be good mimics of CD20. In BLI experiments with a cyclic peptide derived from ECL2 the measured K_d_ for rituximab was only 2 μM, while soluble CD20 and native CD20 bind rituximab with a K_d_ of 2 nM. The similar affinity of soluble CD20 and native CD20 for rituximab suggests that the cleft formed between two copies of ECL2 is the primary structural feature contributing to binding affinity.

The antibody binding studies with rituximab and obinutuzumab indicate that soluble CD20 presents ECL2 in a native-like manner. We also tested binding with the anti-CD20 antibody ofatumumab. The binding epitope for ofatumumab includes residues 71-80 on ECL1 and residues 146-160 on ECL2. Since soluble CD20 does not include ECL1, it is not surprising that only modest affinity was observed for ofatumumab via BLI, and soluble CD20 and ofatumumab failed to form a stable complex when probed with SEC-MAS (Figure S4). This suggest that current engineered soluble CD20 can only be used to discover antibodies or antibody-like molecules that interact with ECL2 only.

In summary, we have shown that computational protein design can be used to present a critical dimer epitope from CD20 in a native-like conformation in a protein that is stable and soluble in water. Furthermore, we demonstrated that soluble CD20 is amenable to yeast display experiments and binds effectively to anti-CD20 binders presented on the surface of yeast. We anticipate that soluble CD20 will be useful for discovering non-antibody binders against CD20 and will be a useful reagent for characterizing newly discovered anti-CD20 antibodies. Our study highlights the use of computational design for epitope scaffolding to create water-soluble analogs of challenging transmembrane proteins, potentially broadening the scope of drug development.

## Materials and Methods

### Rosetta computational design methods for Soluble CD20 Dimer

The cryo-EM structure of full-length CD20 dimer (PDB code: 6VJA) served as the starting point for the design process. We designed chimeras that scaffolded the ECL2 loop and the replaced transmembrane domain with a de novo coiled coil. First, a large family of idealized dimeric coiled-coils with 42-residues were generated. In PyRosetta, we described the superhelical twist and radius of a coiled-coil (Grigoryan and DeGrado 2011a). We then probed which of these coiled coils most closely align structurally (Cα RMSD) with residues from the n-terminal portions of TM4 in the CD20 dimer. More specifically we calculated the root mean square deviation (RMSD) between the Cα atoms on residues 1 and 2 from both chains of the idealized coiled coil with the Cα atoms on residues 184 and 185 (residue numbering from PDB 6VJA) from both chains of CD20. Ten coiled coils, which superimpose with low RMSD (<= 0.25 Å) were selected to build full length models of the designed proteins. These models were constructed by grafting residues 158-185 from CD20 onto the N terminus of the matching coiled coils, resulting in protein chains with sixty-eight residues.

Sequences were designed for the CD20/coiled coil chimeras using the FastDesign protocol in the molecular modeling program Rosetta. FastDesign iterates between rotamer-based sequence optimization and backbone refinement to find low energy sequence/structure pairs (Maguire et al. 2021). During these simulations the sequence of the ECL2 was fixed as the native sequence and the residues in the coiled coil were allowed to vary to the following amino acids ADEIKLNQRSTV. The native disulfide between residues 167 and 183 in the ECL2 was maintained during the simulations. To preserve sequence symmetry between the two chains of the homodimer, symmetry restraints were applied to designable residues. The total energy of the design model as calculated with the full atom energy function (energy function beta_nov15) binding energy normalized by interface area (“dG_separated/dSASA”). From 400 design trajectories (40 for each coiled-coil template), 8 sequences were selected for experimental characterization. The scripts used for the design runs are provided in the supplementary materials (supplementary table 1).

### Cloning, expression, and purification of soluble CD20

DNA encoding the design sequences were synthesized (Twist biology) and cloned into a modified pαH mammalian expression vector. Two different soluble CD20 constructs, soluble CD20^His^ and soluble CD20^Avi^ (supplementary table 2) were produced. Soluble CD20 was expressed in EXPI293F cells (Thermo Fisher Scientific, catalog A14527) with a human serum albumin signal peptide for secretion to the culture medium. Constructs were transiently transfected according to the manufacturer’s protocol with a modified cell density of 3.0 × 106 cells/ml. EXPI293F cells were maintained at 37°C with 8% CO2 at 250 rpm without antibiotics. Enhancers were added according to the manufacture’s protocol 20 hours after transfection. The cell cultures were harvested 96 hours after transfection and the supernatants were separated from the cells by spinning at 3000 rfc (4°C) and passing through 0.4µm filters. The supernatants were then mixed with penta-Ni resin (Marvelgent) under gentle shaking at 4 °C for 30 minutes to allow protein binding. After incubation, resin was washed with 2.5 column volumes (CV) of 10mM imidazole in high salt Tris buffer (1M NaCI, 50mM Tris base in PH 8), 1 column volumes(CV) of 25mM imidazole in high salt tris buffer and 1 CV of 50mM Imidazole in PBS, and elution was performed by 4 steps of 1 CV PBS with 500mM Imidazole.

*Escherichia coli* Shuffle T7 cells (Thermo Fisher Scientific, USA) were used for protein expression due to their high efficiency and compatibility with T7 promoter-driven expression systems. The gene of interest was cloned into the expression vector pET- 28a(+), containing an N-terminal His-tag as described in supplementary table 2 Soluble CD20^bacteria^. Plasmid DNA was transformed into competent Shuffle T7 cells using the heat-shock method. A single colony was picked and grown in 5 mL of LB medium with the Ampicillin overnight at 37°C, with shaking at 250 rpm. The overnight culture was used to inoculate 50ml LB and then 1L of fresh TB culture. Cells were induced with IPTG(final concentration of 0.5 mM) at OD600 of 0.6-0.8. The temperature was then dropped to 16°C and shake at 250rpm overnight. Cells were harvested by centrifugation at 4,000 x g for 30 minutes at 4°C. The cell pellet was frozen and stored at-80 °C. Cells were lysed in lysis buffer (500mM NaCl, 50mM Tris HCl pH 7.5, 10mM Imidazole, PMSF 1mM, Pepstatin 10 µM, Bestatin 1 µM, Leupeptin 10 µM) by sonication on ice (5 seconds on, 5 seconds off, at 70% amplification for 5 minutes).

The lysate was clarified by centrifugation at 12,000 x g for 30 minutes at 4°C. The supernatant was batch bind with penta-Ni resin (Marvelgent) under gentle shaking at 4 °C for 30 minutes to allow protein binding. The column was washed with 20 column volumes of high salt wash buffer (1M NaCl, 50mM Tris HCl pH 7.5, 10mM Imidazole), 2.5 column volume of wash buffer (500mM NaCl, 50mM Tris HCl pH 7.5, 25mM Imidazole) to remove non-specifically bound proteins. The target protein was eluted with elution buffer (500mM NaCl, 50mM Tris HCl pH 7.5, 500mM Imidazole).

### Generation of CD20 peptide and native CD20

The CD20 epitope peptide NIYN(CEPANPSEKNSPSTQYC)YSIQ-K(Biot)-NH2 was synthesized and purified by the UNC High-throughput Peptide Synthesis and Array Facility. Native CD20 was recombinantly expressed and purified by cell membrane solubilization. Native CD20 constructs were synthesized (Twist biology) and cloned into a pFastBac vector. Recombinant baculovirus was generated using the Bac-to-Bac® Baculovirus Expression System (invitrigen). Sf9 cells were infected for protein production and harvested 72 hrs post-infection. The sequence of native CD20 was taken from the cryo-EM structure study (Rougé et al. 2020). Native CD20s were expressed in Sf9 cells that was conducted using a baculovirus encoding for recombinant Multidrug resistance-associated protein 4(MRP4) generated from a pFastBac-his6-TEV-CD20-FLAG-AviTag.

Native CD20 was purified by solubilizing the membranes from a large scale Sf9 cell culture. 2L Sf9 cells were spin down 3200 g for 15–30 min at 4 C and resuspended in 100ml 25 mM Tris pH 7.5, 150 mM NaCl with complete EDTA-free protease-inhibitor cocktail tablets (Roche). The cell suspension was homogenized by 30-40 strokes in a 40 mL Dounce homogenizer on ice, then spun down at 8,000 rpm for 20 min. Supernatant that contains the cell membrane component was separated by spinning at 40,000 rpm at 4 °C for 1 hour. Isolated membrane components were resuspended in buffer with 1% (w/v) GDN (Anatrace) and 0.2% (w/v) cholesterol hemisuccinate in 25 mM Tris pH 7.5, 150 mM NaCl with surfactant as suggested (Rougé et al. 2020). Solubilization was achieved using gentle agitation for at least 4 hours at 4 °C. Ultracentrifugation at 40,000 rpm at 4 °C for 30 min was used to remove insoluble debris. Clarified supernatant was used for 1^st^ Nickle-His purification with penta-Ni resin (Marvelgent). Supernatant were incubated with pre-equilibrated penta-Ni resin under gentle shaking at 4 °C overnight and then washed with 10 column volumes of detergent buffer (25 mM Tris pH 7.5, 150 mM NaCl, 0.02% GDN) plus 10 mM imidazole and eluted with detergent buffer plus 250mM imidazole. The eluted samples were loaded onto a Superdex S200 10/300 GL column (GE Healthcare) pre-equilibrated with detergent buffer, and major fractions were collected for the 2^ed^ TEV cutting purification. These fractions were inculcated with His-tagged TEV protease to cleave the tags, followed by passage through a fresh Ni-NTA column equilibrated with detergent buffer. Untagged CD20 was recovered in the flow-through and wash fractions after washing and elution.The eluted samples were loaded onto a Superdex S200 10/300 GL column (GE Healthcare) pre-equilibrated with detergent buffer, and the major fractions were collected. These fractions were used in 2ed TEV cutting purification. Fractions were incubated overnight with His-tagged TEV protease prepared in-house to cleave the tags and then passed through a new penta-Ni column that equilibrated with detergent buffer. After washing and elution steps, as previously described, the flow-through and wash fractions containing the untagged CD20 were collected.

### Assessment of De Novo CD20 protein stability with Circular dichroism (CD)

Each purified protein was centrifuged at 14,000 × g for 10 min at 4°C and the supernatant diluted to 0.5uM, 2uM, 20uM and 40Um in PBS. Circular dichroism spectra and temperature melts were acquired with a Jasco CD J-815 C. Thermal unfolding was monitored by ramping at 1°C intervals from 20°C to 90°C and measuring the CD signal at 222 nm during the temperature ramp. Molar ellipticity was calculated with the following expression: [θ]=θ/(10 × c × l) where l is the path length in cm. The midpoint for thermal unfolding was determined by fitting the data to the temperature dependent Gibbs-Helmhotz Equation using non-linear regression tools in Microsoft Excel.

### Bio-Layer Interferometry (BLI) measurements for binding affinity

Bio-Layer Interferometry (BLI) assays were performed to determine the binding affinity between selected antibodies and native or de novo designed CD20 receptors. Biotinylated native or soluble CD20 was immobilized on streptavidin SA biosensors (FortéBio / Molecular Devices) at a concentration of 30ug/ml. The biosensors were moved to wells with 5-fold serial dilutions of antibodies to assess association and dissociation. One additional reference well with unloaded biosensors was mixed with the highest antibody concentration well and were used to normalize data. The association and dissociation of 5 concentrations of antibodies were used to fit kinetic data to a 2:1 binding mode to determine the affinity constant (KD) using the association (k_on_) and dissociation (k_off_) rates using software (Octet® Analysis Studio). For de novo CD20, kinetic buffer (kinetics buffer 10X (No. 18-1105) was used for all washing, dilution and sensor activation. For native CD20, PBS buffer containing 0.02% GDN supplemented was use for all washing, dilution and sensor activation.

### Size-Exclusion Chromatography-Multiangle Light Scattering (SEC-MALS)

Purified soluble CD20 and CD20 Fab antibodies were mixed in a molar ratio 1.5:1 and incubated for 30 min at 4°C with rotation to form complexes. Proteins were then injected onto a superdex column (Superdex 200 Increase 10/300 GL Cytiva 28-9909-44) equilibrated in PBS at 0.5ml/min and probed with an inline light scattering detector (Wyatt DAWN HELEOS II static light scattering system).

### Mass photometry

Purified soluble CD20 and CD20 Fab antibodies were mixed in a ratio 1.5:1 and incubated for 30 min at 4°C with rotation to form complexes, and then further purified with gel filtration (Superdex 200 Increase 10/300 GL Cytiva 28-9909-44). The purified complexes were prepared at a concentration of 17 μM in PBS buffer and then diluted to a concentration of 170 nM immediately before mass photometry. 2uL of sample were loaded into a 10 μL buffer-containing well on clean coverslips. The data were acquired using a Refeyn One mass photometer, Refeyn AcquireMP for 60s and analyzed with the DiscoverMP (v2.3) software.

### Yeast surface display

The single-chain variable fragment (scFv) of rituximab was cloned into the pCTCON2 yeast display vector, which includes a myc-tag for expression verification. *Saccharomyces cerevisiae* strain EBY100 competent cell was prepared and transformed with the pCTCON2-scFv plasmid using electroporation following a standard transformation protocol (Chao et al. 2006). Transformed yeast cells were recovered on YPD media and resuspended in SDCAA media. A single colony was picked and cultured twice for 48 hours in 5 mL SDCAA medium at 30°C. 5 × 10^7^ cells were used to inoculate 5 ml SGCAA culture and grown for 20 hours at 20°C, 220 rpm for induction.

Binding assays were performed using three forms of biotinylated CD20 antigens: CD20 peptide (10 µM), native CD20 (0.6 µM), and a serial dilution of soluble CD20 ranging from 0.8 nM to 0.6 µM. The binding signal of CD20 antigens were detected through Alexa Fluor™ 647-conjugated streptavidin (Invitrogen, S21374) that binds to the biotinylated CD20 antigens. Surface expression of the scFv was confirmed by flow cytometry using FITC-conjugated anti-Myc tag antibody (Abcam, ab1394). After induction, 5 × 10^6^ yeast cells displaying rituximab scFv were collected and were washed twice with PBSF buffer (137 mM NaCl, 2.7 mM KCl, 8 mM Na2HPO4, and 2 mM KH2PO4, supplemented with 0.1% BSA). Cells were incubated with FITC-conjugated anti-Myc tag antibody for 1 hour at 4°C with shaking, followed by three washes with PBSF buffer. CD20 antigens were preloaded with Alexa Fluor™ 647-conjugated streptavidin at a molar ratio of 4:1 for 30 minutes at room temperature before being incubated with yeast cells for 1 hour at 4°C with shaking. After incubation, cells were washed three times with PBSF buffer.

Flow cytometry was performed on a Thermo Fisher Attune NxT instrument. The expression signal (myc-positive) and the antigen binding signal was detected in the FITC channel and Alexa Fluor™ 647 channel respectively. Control samples, that labeled with fluorescent Streptavidin conjugate but without antigen, were used to establish background fluorescence levels. Flow cytometry data were analyzed using FlowJo software. The binding positive rate of each CD20 antigens were quantified as the percentage of double-positive cells (scFv-expressing and antigen-binding cells) among total scFv-expressing cells.

## Supporting information

protein design scripts

## Abbreviations

CMPs: Complex membrane proteins
ECDs: Extracellular Domains
CDC: Complement-Dependent Cytotoxicity
PCD: Programmed Cell Death
mAbs: Monoclonal Antibodies
Fab: fragment antigen-binding
scFv: single chain variable fragment
CD: Circular Dichroism
Tm: Melting Temperature
TM: transmembrane helix
SEC-MALS: Size Exclusion Chromatography-Multi Angle Light Scattering
BLI: Bio-Layer Interferometry
SPR: Surface Plasmon Resonance
RTX: Rituximab
OBZ: Obinutuzumab
RMSD: Root-Mean-Square Deviation

## Acknowledgements

The authors thank Kiara Thompson for help in expressing soluble CD20 in *e.coli*, Nicole Hajicek and Runfan Yang in John Sondek’s lab for advice regarding membrane protein expression in SF9 cells, and Matthew Cummins for advice and discussion of the project. We are grateful to Julia Koehler Leman for her help in manuscript writing. We thank the UNC Macromolecular Interactions Facility and UNC Flow Cytometry Core Facility for advice and instrumentation. This work was supported by the NIH grant R35GM131923 (BK).

## Conflict of interest statement

The authors have no conflict of interests related to this research.

## Data availability statement

The data that support the findings of this study are available from the corresponding author upon request.

## Notes

### Competing Interest Statement

The authors have declared no competing interest.

