## Supplementary figures and tables for "Design of a Water-Soluble CD20 Antigen with Computational Epitope Scaffolding"

Table of contents:

**Supplementary table 1.** Pyrosetta and Rosetta protocols used to design soluble CD20.

**Supplementary table 2.** List of designed soluble and native CD20 protein sequences and constructs.

**Supplementary table 3.** List of anti-CD20 antibodies used in soluble expression and yeast display.

**Supplementary Figure 1.** Expression of all designed soluble CD20 sequences and native CD20

**Supplementary Figure 2.** Expression and purification of Native CD20 with membrane solubilization

**Supplementary Figure 3.** Components of distinct peaks from size exclusion chromatography with soluble CD20 and Fab antibody mixtures.

**Supplementary Figure 4.** Binding affinity and binding complex characterization of ofatumumab Fab and soluble CD20.

**Supplementary Table 1. Pyrosetta and Rosetta protocol for soluble CD20 design.** Soluble CD20 were designed by scaffold generation and sequence design with PyRosetta and Rosetta respectively. Scripts, input files and example outputs are included in the supplementary file cd20_scripts.zip.

| Design stage | scripts | Input | Example Output |
|---|---|---|---|
| PyRosetta Scaffold generation | `helix_match_cd20.py` | Partial structure from PDB 6VJA: `cd20_frag.pdb` | Score file: `rnd1_scorefile.txt`<br>Coil-coil α helix dimer: `rnd1_2_4.9_-14.0_-4.2_cc.pdb`<br>CD20 epitope fragment: `rnd1_2_4.9_-14.0_-4.2_cd20.pdb`<br>Soluble CD20 scaffold: `rnd1_2_4.9_-14.0_-4.2_chimera.pdb` |
| Rosetta Sequence design | `design_cd20_cc.xml`<br>`run.sh`<br>`C2_Z.sym`<br>`Option`<br>`resfile_cd20_cc` | Scaffold input: `rnd1_28_5.1_-5.5_-4.2_chimera.pdb` | Score file: `scorefile.sc`<br>Designed soluble CD20 model: `rnd1_28_5.1_-5.5_-4.2_chimera_design_0001.pdb` |

**Supplementary table 2.** Protein sequences and constructs.

| Protein | | Construct | Sequence |
|---|---|---|---|
| Cd20 peptide | | | NIYN(CEPANPSEKNSPSTQYC)YSIQ-K(Biot)-NH2 |
| Native CD20 | Expression in SF9 | -his6-TEV-nativeCD20-FLAG-**AviTag** (25.1kDa) | MGHHHHHHGSGENLYFQSQAMRESKTLGAVQIMNGLFHIALGGLLMIPAGIYAPICVTVWYPLWGGIMYIISGSLLAATEKNSRKCLVKGKMIMNSLSLFAAISGMILSIMDILNIKISHFLKMESLNFIRAHTPYINIYNCEPANPSEKNSPSTQYCYSIQSLFLGILSVMLIFAFFQELVIAGIVENEWKRGSSGGSDYKDDDDKSSG**GLNDIFEAQKIEWHE** |
| Soluble CD20 | Soluble CD20$^{His}$ | -HSA-his8-**TEV**-solubleCD20-(10.4kDa) | MKWVTFISLLFLFSSAYSHHHHHHHHGGGSGGGS**ENLYFQS**HTPYINIYNCEPANPSEKNSPSTQYCYLAQELLSKNRHLENEVKRLKKLVDDLEDELKAQKEKYKAIS |
| | Soluble CD20$^{Avi}$ | -HSA-solubleCD20-FLAG-his8-**AviTag**-(14.9kDa) | MKWVTFISLLFLFSSAYSHTPYINIYNCEPANPSEKNSPSTQYCYLAQELLSKNRHLENEVKRLKKLVDDLEDELKAQKEKYKAISGSSGGSDYKDDDDKSSGHHHHHHHHSSG**GLNDIFEAQKIEWHE** |
| | Soluble CD20$^{bacteria}$ | -solubleCD20-**TEV**-his8-**AviTag**- (12.3kDa) | MGHTPYINIYNCEPANPSEKNSPSTQYCYLAQELLSKNRHLENEVKRLKKLVDDLEDELKAQKEKYKAISSG**ENLYFQS**SSGHHHHHHSSGG**LNDIFEAQKIEWHE** |

**Supplementary table 3.** Sequences of anti-CD20 antibodies

| Protein | Construct | Sequence |
|---|---|---|
| RTX Fab | Heavy chain: -<br>Fab_H-his8<br>Light chain:<br>Fab_L<br>(48.9kDa) | H:<br>QVQLQQPGAELVKPGASVKMSCKASGYTFTSYNMHWVKQTPG<br>RGLEWIGAIYPGNGDTSYNQKFKGKATLTADKSSSTAYMQLS<br>SLTSEDSAVYYCARSTYYGGDWYFNVWGAGTTVTVSAASTKG<br>PSVFPLAPSSKSTSGGTAALGCLVKDYFPEPVTVSWNSGALT<br>SGVHTFPAVLQSSGLYSLSSVVTVPSSSLGTQTYICNVNHKP<br>SNTKVDKKVEPKSCENLYFQSSSGHHHHHHHH<br>L:<br>QIVLSQSPAILSASPGEKVTMTCRASSSVSYIHWFQQKPGSS<br>PKPWIYATSNLASGVPVRFSGSGSGTSYSLTISRVEAEDAAT<br>YYCQQWTSNPPTFGGGTKLEIKRTVAAPSVFIFPPSDEQLKS<br>GTASVVCLLNNFYPREAKVQWKVDNALQSGNSQESVTEQDSK<br>DSTYSLSSTLTLSKADYEKHKVYACEVTHQGLSSPVTKSFNR<br>GEC |
| OBN Fab | Heavy chain: -<br>Fab_H-his8<br>Light chain:<br>Fab_L<br>(49.8kDa) | H:<br>QVQLVQSGAEVKKPGSSVKVSCKASGYAFSYSWINWVRQAPG<br>QGLEWMGRIFPGDGDTDYNGKFKGRVTITADKSTSTAYMELS<br>SLRSEDTAVYYCARNVFDGYWLVYWGQGTLVTVSSASTKGPS<br>VFPLAPSSKSTSGGTAALGCLVKDYFPEPVTVSWNSGALTSG<br>VHTFPAVLQSSGLYSLSSVVTVPSSSLGTQTYICNVNHKPSN<br>TKVDKKVEPKSCENLYFQSSSGHHHHHHHH<br>L:<br>DIVMTQTPLSLPVTPGEPASISCRSSKSLLHSNGITYLYWYL<br>QKPGQSPQLLIYQMSNLVSGVPDRFSGSGSGTDFTLKISRVE<br>AEDVGVYYCAQNLELPYTFGGGTKVEIKRTVAAPSVFIFPPS<br>DEQLKSGTASVVCLLNNFYPREAKVQWKVDNALQSGNSQESV<br>TEQDSKDSTYSLSSTLTLSKADYEKHKVYACEVTHQGLSSPV<br>TKSFNRGEC |
| OFA Fab | Heavy chain: -<br>Fab_H-his8<br>Light chain:<br>Fab_L<br>(49.6kDa) | H:<br>QVQLQQPGAELVKPGASVKMSCKASGYTFTSYNMHWVKQTPG<br>RGLEWIGAIYPGNGDTSYNQKFKGKATLTADKSSSTAYMQLS<br>SLTSEDSAVYYCARSTYYGGDWYFNVWGAGTTVTVSAASTKG<br>PSVFPLAPSSKSTSGGTAALGCLVKDYFPEPVTVSWNSGALT<br>SGVHTFPAVLQSSGLYSLSSVVTVPSSSLGTQTYICNVNHKP<br>SNTKVDKKVEPKSCENLYFQSSSGHHHHHHHH<br>L:<br>QIVLSQSPAILSASPGEKVTMTCRASSSVSYIHWFQQKPGSS<br>PKPWIYATSNLASGVPVRFSGSGSGTSYSLTISRVEAEDAAT<br>YYCQQWTSNPPTFGGGTKLEIKRTVAAPSVFIFPPSDEQLKS<br>GTASVVCLLNNFYPREAKVQWKVDNALQSGNSQESVTEQDSK<br>DSTYSLSSTLTLSKADYEKHKVYACEVTHQGLSSPVTKSFNR<br>GEC |
| RTX scFv for yeast display | -linker-scFv-<br>linker-cMyc- | GGGGSGGGGSGGGGSASQIVLSQSPAILSASPGEKVTMTCRA<br>SSSVSYIHWFQQKPGSSPKPWIYATSNLASGVPVRFSGSGSG<br>TSYSLTISRVEAEDAATYYCQQWTSNPPTFGGGTKLEIKGGG<br>GSGGGGSGGGGSQVQLQQPGAELVKPGASVKMSCKASGYTFT<br>SYNMHWVKQTPGRGLEWIGAIYPGNGDTSYNQKFKGKATLTA<br>DKSSSTAYMQLSSLTSEDSAVYYCARSTYYGGDWYFNVWGAG<br>TTVTVSGSGSGSGSGSEQKLISEEDL |

a.

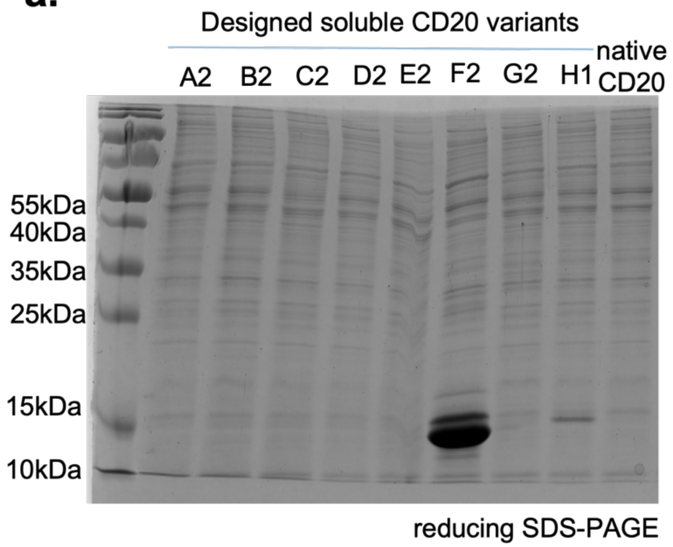


**Supplementary Figure 1. Expression of designed soluble CD20 sequences and native CD20.** 8 designed soluble CD20 variants (named A2 to H1) and native CD20 were produced using a modified mammalian expression vector and expressed as secreted proteins in Expi293 cells and purified with metal affinity chromatography. Designed soluble CD20 variants F2 and H1 showed detectable amount of protein. Native CD20 cannot be secreted as a soluble protein from Expi293 cells. Reducing SDS-page (12% acrylamide) gel were used. Design F2 was renamed “soluble CD20”.

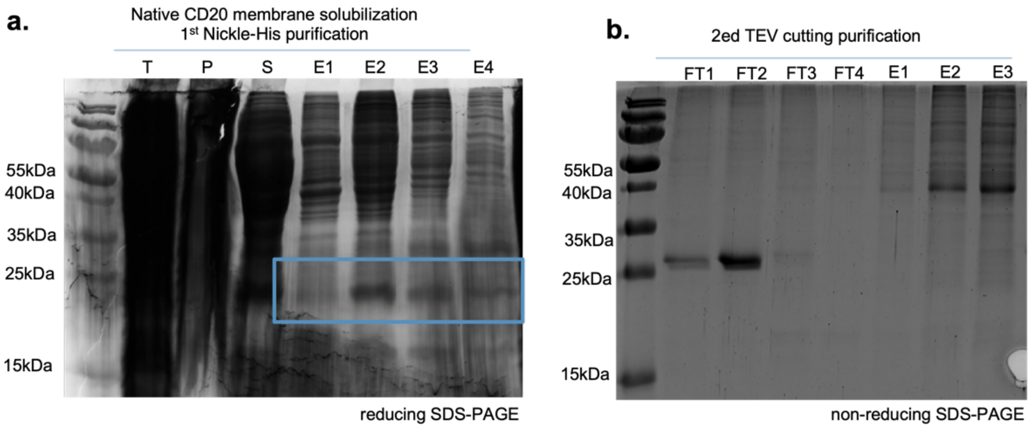


**Supplementary Figure 2. Expression and purification of Native CD20 with membrane solubilization.** Native CD20 was recombinantly expressed and purified with two steps through cell membrane solubilization. (a) In 1st Nickle-his purification, the native CD20 at around 25.1 kDa were collected through membrane solubilization. The total lysate "T", pallet insoluble fraction obtained after membrane solubilization "P", supernatant of membrane fraction "S", and 4 elution samples "E1, E2, E3,E4". (b) After cleavage with TEV protease, native CD20 was collected in the flow through (FT1 and FT2) to yield the final purified product.

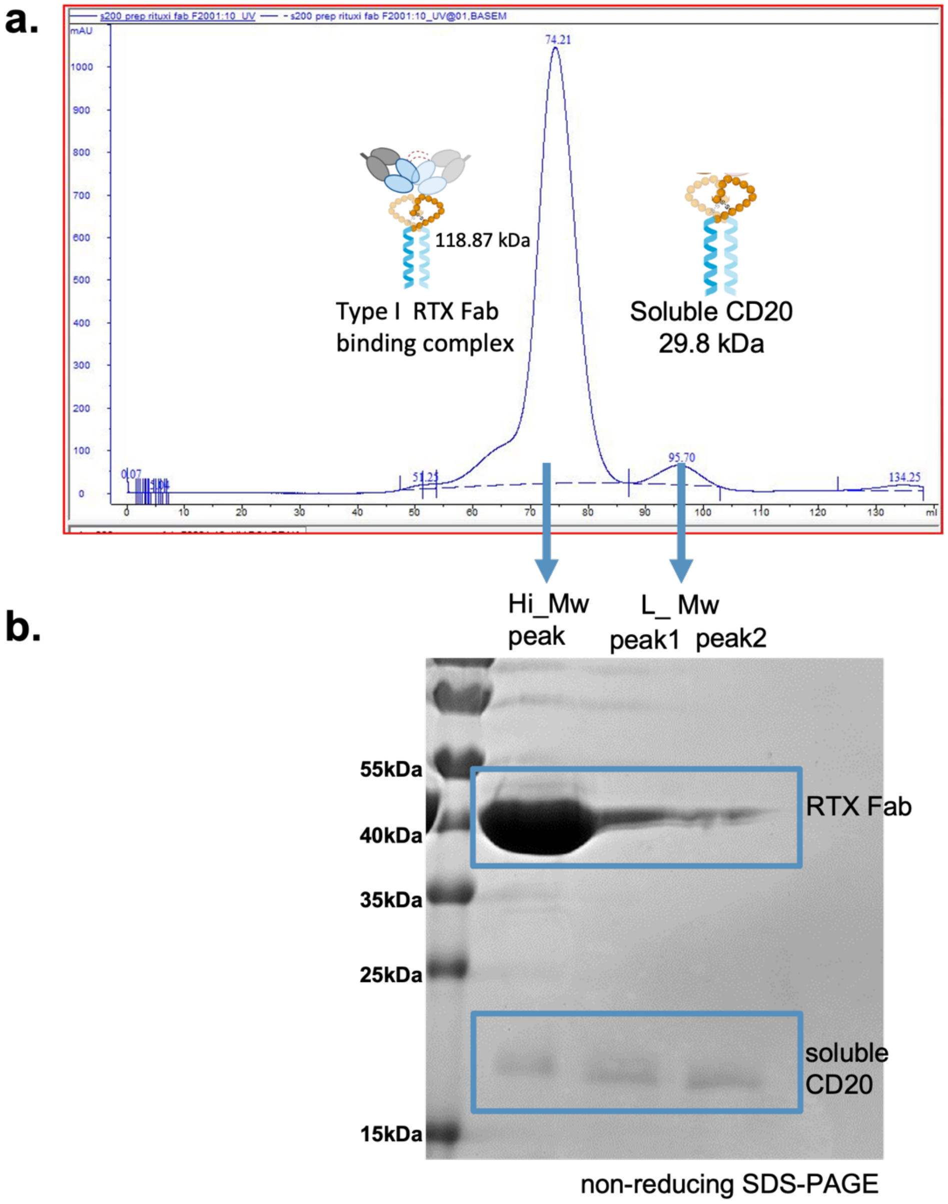


**Supplementary Figure 3.** RTX Fab and soluble CD20 co-elute during size exclusion chromatography (SEC). (a) SEC analysis of the interaction between RTX Fab and soluble CD20. The major peak (Hi_Mw) at 118.87 kDa corresponds to the RTX Fab-soluble CD20 binding complex. A separate peak (L_Mw) at 29.8 kDa represents unbound soluble CD20, which appears when the components are mixed in a molar ratio of 2:2.5 (b) SDS-PAGE analysis of the SEC fractions confirms the formation of the RTX Fab-soluble CD20 complex. The upper band corresponds to the RTX Fab, and the lower band corresponds to soluble CD20. These results demonstrate that soluble CD20 forms a stable complex with RTX Fab, with excess soluble CD20 appearing as a distinct unbound fraction.

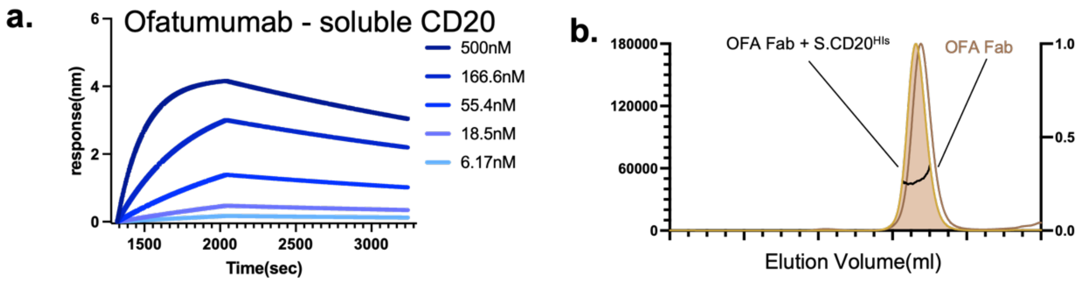


**Supplementary Figure 4.** Biophysical characterization of the interaction between ofatumumab Fab and soluble CD20. (a) Bio-layer interferometry (BLI) sensor grams showing the binding of ofatumumab Fab to soluble CD20.The binding affinity of soluble CD20 to type I antibody ofatumumab is 24.1 nM. (b) SEC-MALS analysis of the soluble CD20-Ofatumumab (OFA) Fab mixture. No stable complex formation was detected compared to OFA fab control. While OFA shows decent binding affinity on soluble CD20 with ECL2 only, it cannot form binding complex with soluble CD20 lacks OFA epitope residues on ECL1.
