## Supplementary material for "Design of a Water-Soluble CD20 Antigen with Computational Epitope Scaffolding": protein design scripts: helix_match_cd20.py

```
from pyrosetta import *
from pyrosetta.teaching import *
import pyrosetta.rosetta.protocols.helical_bundle as helical_bundle
import pyrosetta.rosetta.core.simple_metrics.per_residue_metrics as per_residue_metrics
import numpy as np
import itertools

init(extra_options = "-ex1 -ex2 -extrachi_cutoff 1")

helix_length = 42
output_counter = 0
output_tag = "rnd1"    # 


## coiled coil range to sample
r0_array = np.linspace(4.9,5.1,3)
delta_omega1_array = np.linspace(-18,-2,5)   # degrees, helix 1
#omega0_array = np.linspace(-4.207,-4.207,1)    # degrees, superhelical twist
omega0_array = np.linspace(-4.2,-4.2,1)    # degrees, superhelical twist
#omega1 = np.radians(2*360/7)  #  radians, twist per residue in a_helix, determines # of layer

## residues to match
cd20_match = [27,28,57,58]
cc_match = [1,2,1+helix_length,2+helix_length]


pose_cd20_in = pose_from_file("cd20_frag.pdb")

################

scf_name ='%s_%s'%(output_tag,"scorefile.txt")
scorefile = open(scf_name,"w")
l = len(output_tag)
header = str.ljust("file_name",l+11)  + "r0 d_omeg1  omeg0  rmsd \n"  
scorefile.write(header)

pose_cc = Pose()

for r0,delta_omega1,omega0 in itertools.product(r0_array,delta_omega1_array,omega0_array):
        pose_cd20 = Pose()
        pose_cd20.assign(pose_cd20_in)
        output_counter = output_counter + 1
        
        mb_mover = helical_bundle.MakeBundle()
        mb_mover.set_use_degrees()
        mb_mover.set_default_helix_length(helix_length)
        mb_mover.set_symmetry(2)
        mb_calc = mb_mover.default_calculator_nonconst()
        mb_calc.parameter(helical_bundle.BPC_set_bondangles).set_value(True)
        mb_calc.parameter(helical_bundle.BPC_set_bondlengths).set_value(True)

        # helix parameters
        mb_calc.parameter(helical_bundle.BPC_r0).set_value(r0)
        mb_calc.parameter(helical_bundle.BPC_delta_omega1).set_value(delta_omega1)
        mb_calc.parameter(helical_bundle.BPC_omega0).set_value(omega0)
        mb_calc.parameter(helical_bundle.BPC_invert_helix).set_value(False)  # 
        mb_calc.parameter(helical_bundle.BPC_delta_omega0).set_value(0.)  # 
        mb_calc.parameter(helical_bundle.BPC_z1_offset).set_value(0.)  # 
        mb_mover.add_helix()
        
        mb_mover.apply(pose_cc)
        name='%s_%s_%s_%s_%s'%(output_tag,output_counter,r0,delta_omega1,omega0)
        #name=output_tag+"_"+str(output_counter)+".zfill(6)

        atom_map = rosetta.core.id.AtomID_Map_AtomID()
        rosetta.core.pose.initialize_atomid_map( atom_map, pose_cd20, rosetta.core.id.AtomID.BOGUS_ATOM_ID() )

        for i in range(len(cd20_match)):
            atom_cc = rosetta.core.id.AtomID(2,cc_match[i])   # atom 2 is the Calpha
            atom_cd20 = rosetta.core.id.AtomID(2,cd20_match[i])
            atom_map.set(atom_cd20,atom_cc)

#atid = lambda x : rosetta.core.id.AtomID(2,x)
#mapping = {atid(cc_list[i]):atid(cd20_list[i]) for i in range(4)}

        rmsd = rosetta.core.scoring.superimpose_pose(pose_cd20,pose_cc,atom_map)
        print("rmsd =", rmsd)

        scorefile_line='%s %5.1f %6.1f %6.1f %6.2f'%(name,r0,delta_omega1,omega0,rmsd)
        scorefile.write(scorefile_line+"\n")

        # for top hits create chimeric pose, output just one chain so that it can be used in symmetric mode design
        if rmsd < 0.26:
            pose_cd20.dump_pdb(name+'_cd20.pdb')
            pose_cc.dump_pdb(name+'_cc.pdb')

            pose_cd20.delete_residue_range_slow(28,61)    #deleting end of c-terminal helix and other chain
            pose_cc.delete_residue_range_slow(helix_length+1,2*helix_length)  # other chain
            pose_cc.delete_residue_range_slow(1,1) # deleting first residue
            rosetta.core.pose.append_pose_to_pose(pose_cd20,pose_cc,False)
            ft = FoldTree()
            ft.add_edge(1,27+helix_length-1,-1)
            pose_cd20.fold_tree(ft)   # append_pose_to_pose uses jumps to connnect poses, we want one chain
            pose_cd20.dump_pdb(name+'_chimera.pdb')
        
scorefile.close()
#pose_cd20.dump_pdb("cd20_match.pdb")
#pose_cc.dump_pdb("cc_match.pdb")
```
