## Supplementary material for "Design of a Water-Soluble CD20 Antigen with Computational Epitope Scaffolding": protein design scripts: run.sh

```
#!/bin/bash
#SBATCH --mail-user=
# Email address
#SBATCH --mail-type=ALL
#SBATCH --time=240
#SBATCH --tasks-per-node=44
#SBATCH --ntasks=22
#SBATCH --distribution=cyclic:cyclic
#SBATCH -o log.txt
#SBATCH -e error.txt
#SBATCH -p debug_queue

####### ####### ####### ####### SUBMISSION COMMAND
$MPI_HOME/bin/mpirun /nas/longleaf/home/zyyao/Rosetta/main/source/bin/rosetta_scripts.mpiserialization.linuxgccrelease -parser:protocol design_cd20_cc.xml -s rnd1_28_5.1_-5.5_-4.2_chimera.pdb -scorefile scorefile.sc -out:suffix _design -mpi_tracer_to_file mpi -overwrite @option
```
